# E156G and Arg158, Phe-157/del mutation in NTD of spike protein in B.1.617.2 lineage of SARS-CoV-2 leads to immune evasion through antibody escape

**DOI:** 10.1101/2021.06.07.447321

**Authors:** Armi M Chaudhari, Dinesh Kumar, Madhvi Joshi, Amrutlal Patel, Chaitanya Joshi

## Abstract

Emerging variants of SARS-CoV-2 with better immune escape mechanisms and higher transmissibility remains a persistent threat across the globe. B.1.617.2 (Delta) variant was first emerged from Maharashtra, India in December, 2020. This variant is classified to be a major cause and concern of the second wave of COVID-19 in India. In the present study, we explored the genomic and structural basis of this variant through computational analysis, protein modelling and molecular dynamics (MD) simulations approach. B.1.617.2 variant carried E156G and Arg158, Phe-157/del mutations in NTD of spike protein. These mutations in N-terminal domain (NTD) of spike protein of B.1.617.2 variant revealed more rigidity and reduced flexibility compared to spike protein of Wuhan isolate. Further, docking and MD simulation study with 4A8 monoclonal antibody which was reported to bind NTD of spike protein suggested reduced binding of B.1.617.2 spike protein compared to that of spike protein of Wuhan isolate. The results of the present study demonstrate the possible case of immune escape and thereby fitness advantage of the new variant and further warrants demonstration through experimental evidence. Our study identified the probable mechanism through which B.1.617.2 variant is more pathogenically evolved with higher transmissibility as compared to the wild-type.

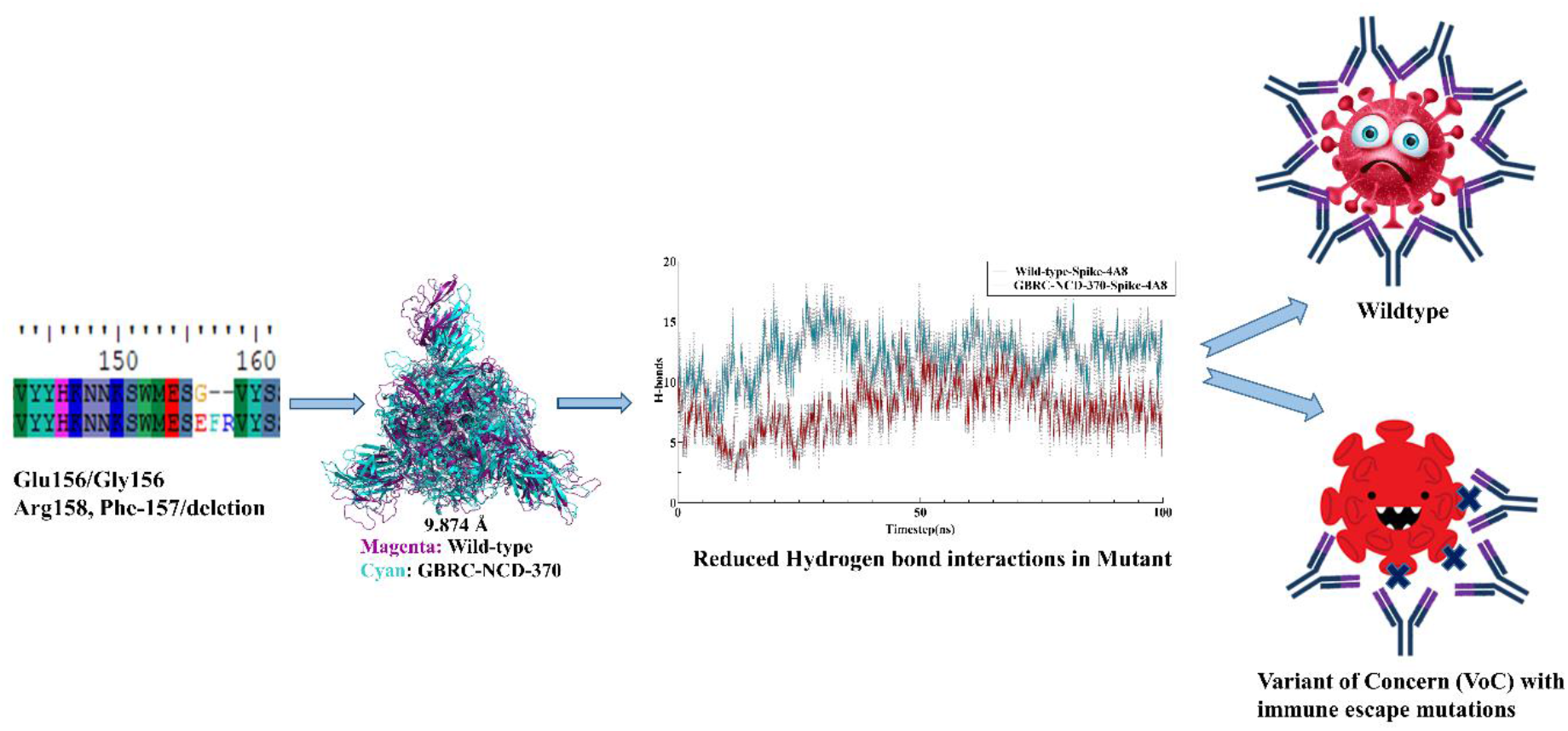

## 1. Introduction

India is witnessing the peak of another COVID-19 wave with over 0.32 million causalities since 2020, and more than 27.4 million confirmed positive cases as per WHO reports accessed on 27^th^ May, 2021. Genome surveillance is a powerful tool to study the viral genomic profile, variants of concern and their epidemiological significance in disease outbreak outcome of the patients. All coronaviruses are positive-sense RNA viruses belonging to the order *Nidovirales* and family *Coronaviridae*. They are characterized by crown-like spikes on their surfaces and large enveloped genome of ∼30 kilobases size. The SARS-CoV-2 genome contains four major structural proteins: spike (S), membrane (M), envelope (E), and nucleocapsid (N) protein. The spike (S) protein mediates entry and attachment of the coronavirus to the host cell surface receptors resulting in fusion and viral entry in the hosts. The membrane (M) protein defines the shape of the viral envelope while the envelope (E) protein and nucleocapsid (N) protein participates in viral assembly and budding of the virion complex in the infected cells [1,2]. SARS-CoV-2 uses ACE2 receptor for host cell entry and the spike protein of SARS-CoV-2 is primed by TMPRSS2 while the role of several other host receptors is partially explored with limited information that may determine the altered virulence and pathogenicity of the evolving SARS-CoV-2 lineages around the globe. SARS-CoV-2 possesses highly efficient and evolved strategies for proteolytic activation of spike, and host proteases have been shown to proteolytically process the spike protein. These include, but are not limited to, endosomal cathepsins, cell surface trans-membrane protease/serine (TMPRSS) proteases, furin, and trypsin that are critical determinants of the virus entry and pathogenesis in humans [3,4]. SARS-CoV-2, in comparison to SARS-CoV, contains a polybasic sequence motif, Arg-Arg-Ala-Arg (RRAR), at the S1/S2 boundary, furin-type cleavage site in its spike protein, which when cleaved can bind and activate neuropilin receptors. Further, research studies indicate that NRP1 enhances SARS-CoV-2 infectivity and is highly expressed in respiratory and olfactory epithelium [5].

Under the prevailing circumstances, the immune response of the patients plays a significant role in determining the disease fighting ability of the body. A myriad of various cell types such as macrophages, alveolar epithelial cells, lymphoid cells, and dendritic cells (DCs) have a major role in the first line of defense. Once the immune system is triggered for the entry of foreign viral pathogens inside the body and that breached the first lines of defense system, several specific molecular and inter-cellular signaling cascades ensure the establishment of the body’s immune response [6,7]. When the invading respiratory viruses evolve mechanism that either circumvent or suppress the innate immune responses to create a window of opportunity for efficient virus replication, thereby often causing disease. The affected innate immune response also impacts subsequent adaptive immune responses, and therefore viral innate immune evasion often undermines fully protective immunity such as lack of virus neutralizing antibodies [8–11]. Further, genetic makeup and evolution in virus also enable them to develop mutations that can cause immune escape and immune evasion in the hosts, thereby increasing the chances of severity and virulence of the pathogenic variants [12]. These variants, which are observed with features such as higher transmittability are categorized as Variants of Concern (VoC) by Public Health England (PHE), UK; CDC, USA, and World Health Organization (WHO) based on their risk assessment criteria of infection severity, susceptibility geographical prevalence, and transmission in humans. Therefore, genomic surveillance studies are essential in monitoring these variants that may even arise in the future pandemics.

New variants of SARS-CoV-2 are emerging challenge for scientific community and public health system in the different geographical regions across the globe. These variants, have been designated as Variant of Concern (VoCs) which has noticeable higher transmissibility and probably more virulent compared to other variants. Genomic structure of such variants suggests that they were evolved to escape the immune system of the host thereby giving them the fitness advantage and thus increased spread among the population. Further, research is needed to establish the mechanism of escape and potential host genetic factor that might help in these evolved pathogenic viral strains of SARS-CoV-2.

Furthermore, understanding of the role of virus-host interactions and immune response during these SARS-CoV-2 infections will be pivotal to ultimately meet these evolving challenges. Eventually, efficacy of the combined innate and adaptive responses is on the host’s side, while the virulence and its capacity to evade the host’s immune responses is on the virus’ side, together, the balance between them dictate the disease outcome in the context of the host-virus interactions. Recent studies on spike protein interactions with monoclonal antibodies 4A8 suggests that N-terminal domain is essential binding site for 4A8 [13]. Some prominent mutations (>99.7% frequency) in virus favors the virus like D614G [14] and some favors the host like C241T [15]. To find the same, this research focus on the mutations in N-terminal domain in B.1.617.2 (now delta) lineage of SARS-CoV-2 and its impact on protein structural changes and antibodies binding using molecular modeling and dynamics approach.

## 2. MATERIAL AND METHODS

### 2.1 Protein complexes used for this study

Variants of Spike protein from SARS-CoV-2 were taken into study. Mutated spike from SARS-CoV-2 used in this study were derived from amino acid sequence submitted in GAISAD with accession number EPI_ISL_2001211. Reference protein with PDB id 7KRQ was used for homology modelling.

### 2.2 Protein modelling and Molecular dynamics simulations

Homology modelling panel implemented in Schrodinger suite release 2021-1 was used to build mutated spike protein with reference protein 7KRQ. Sequence was imported and homology blast search was performed. Crystal structure of 7KRQ was imported in to maestro and protein complex refinement was performed using protein preparation wizard [16]. Missing side chains were added through PRIME and pKa refinement was done with epik [17]. Molecular dynamics simulation for spike protein and Spike-antibody complexes were performed in Schrodinger suite 2021-1 implemented DESMOND till 100 nanoseconds (ns) [18]. Protein structures were refined using OPLS4 force field and altered hydrogen bonds were refilled using structure refinement panel implemented in Schrodinger[19][20]. Particle mesh Ewald method is applied to calculate long rage electrostatic interactions. [21]. The trajectories were recorded at every 1.2 ps intervals for the analysis. TIP3P water molecules were added and 1.5 M Salt concentration was added to neutralize the system [22]. The Martyna–Tuckerman–Klein chain coupling scheme with a coupling constant of 2.0 ps was used for the pressure control and the Nosé–Hoover chain coupling scheme for temperature control [23]. MD simulations were performed at 310.3K temperature. The behaviour and interactions between the protein and protein were analyzed using the Simulation Interaction Diagram tool implemented in Desmond MD package. The stability of complex was monitored by examining the RMSD of the protein and protein atom positions in time. PYMOL was used for obtaining high resolution images [24]. Protein modelling and Molecular dynamics simulations were performed into duplicates.

### 2.3 Molecular docking of Spike protein with monoclonal antibodies using spike and affinity prediction

Variants of spike proteins Wildtype (7KRQ) and Mutant (GBRC-NDC-370) were docked with monoclonal antibody 4A8. Protein structures were prepared using protein-preparation wizard [16]. After structure refinement of protein, PIPER was used for the protein-protein docking [25]. For binding residues (as shown in figure 2A) detection among both receptor (spike) and ligand (antibody-4A8) attraction forces were applied with <3Å cut off. 70000 docking poses were checked for fulfilling the criteria of distance restrains applied for the binding sites residues. Recently deposited crystal structure of spike protein binding with monoclonal antibody was taken for applying the restraint file showing list of spike residues binding with residues of 4A8. Top 30 poses were generated and a pose with highest free glide energy was used for the MD analysis.

Alanine residue scanning was performed for the binding affinity prediction in PDB deposited spike antibody complex with id 7C2L [13]. Binding site residues were mutated to alanine in order to bind the pivotal residues involved into direct binding with antibody. Positive value of Δ affinity indicated that while mutating binding sites residues to alanine, binding is hindered due to small side chains of alanine and which in terms implies that those important residues were essential for direct affinity with antibody [26,27]. Residue alanine scanning panel present in Biologics of Schrodinger 2021-1 is used to perform the above task.

### 2.4 Binding energy calculation

Binding energy for protein-protein complex was calculated in the form of Prime Molecular Mechanics-Generalized Born Surface Area (MMGBSA) using thermal_mmgbsa.py implemented in PRIME module of Schrodinger [28–30]. ΔG of protein-protein complex was calculated using following equation.

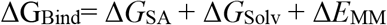

VSEB solvation model and OPLS4 force-field were used for calculation of MMGBSA. Protein-protein complex system seems to have stable RMSD pattern after 60ns. These frames were used to calculate MMGBSA. First energy minimized structure out of 30 was used to find dominant interacting residues among spike. Interaction image was taken in new version of Schrodinger 2021-2 where protein-protein interaction images can be taken in Biologics.

### 2.5 Dynamics cross-correlation matrix (DCCM) and Principal Component analysis (PCA)

Correlative and anti-correlative motions play a vital role in recognition and binding in a biological-complex system which can be prevailed by MD simulation trajectory by determining the covariance matrix about atomic fluctuations [31]. The extent of correlative motion of two residues (or two atoms or proteins) can be symbolized by the cross-correlation coefficient, C_ij_. It is defined by following equation:

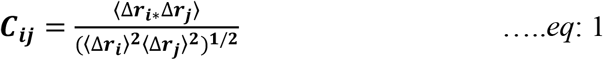

From above equation, i (j) explains ith (jth) two residues (or two atoms or proteins), Δri (Δrj) is the displacement vector corresponding to ith (jth) two residues (or two atoms or proteins), and ⟨. . ⟩ stand for the ensemble average. The value of C_ij_ is from 1 to −1. +C_ij_ implies positively correlated movement (the same direction) indicated into blue color, and −C_ij_ implies anti-correlated movement (opposite direction) indicated into red color. The higher the absolute value of C_ij_ is, the more correlated (or anti-correlated) the two residues (or two atoms or proteins).

PCA is an implicit tool to unsheathe the essential information from MD trajectories by pulling out global slow motions from local fast motions[32]. To perform PCA, the covariance matrix C was calculated initially. The elements C_ij_ in the matrix C are defined as:

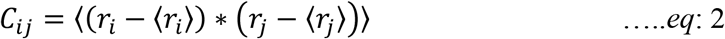

From equation 2, r_i_ and r_j_ are the instant coordinates of the i^th^ or j^th^ atom, ⟨*r_i_*⟩ and ⟨*r_j_*⟩ and mean the average coordinate of the i^th^ or j^th^ atom over the ensemble. The principal components (PCs) were calculated by diagonalization and obtaining the eigenvectors and eigenvalues for the covariance matric C. The principal components (PCs) are the projections of a trajectory on the principal modes, of which usually the first few ones are largely responsible for the most important motions. DCCM and PCA both were analyzed using Schrodinger 2021-1 implemented python script *run trj_essential_dynamics.py* script of Desmond [18].

## 3. Result and discussion

Spike protein of SARS-CoV-2 is known to bind ACE-2 receptor mediating virus entry. Spike protein is more prone to mutations. For better penetrance viral spike had gone through several mutations like D614G for increasing spike density and infectivity, E484K for reducing the antibody neutralization, N501Y and K417N for altering spike interacting with human ACE and human derived antibody [33–35]. Our study focuses on major deletion occurred in NTD of spike protein at nucleotide position 22029-22035 (6bp) which in-terms induce 2 amino-acid deletions Arg158, Phe-157/del and one amino acid mutation E156G.

### 3.1 Mutational landscape of spike protein and its effects with respect to B.1.617.2 lineage

Mutations listed in table 1, were impacting major structural change or not were studied through structural alignment of both variants of spike proteins in Pymol. Superimposed structure of wild-type and GBRC-NCD-370 spike with alignment RMSD 6.905 Å was shown in figure 1A. RMSD value higher than 1Å suggests that these mutations were imparting significant structural change in both variants of spike [36]. Among the list of overall mutations, unique mutations E156G and Arg158, Phe-157/del were falling in NTD of spike. Figure 1D, explains the effect of these mutations in changing amino acids conformation in ball and stick form. One can visualize the difference in alignment of amino acids in NTD within both variants which in terms effects the change in intermolecular contacts within spike (Figure 1E). Due to these mutations intra-atomic contacts within amino acids of mutant (GBRC-NCD-370) is drastically increased with respect to the wild-type. In figure 1D, these mutations change intra-atomic contacts as such that in wild-type Glu-156 is interacting with Phe-140 and Arg158 by forming only one hydrogen bond and other hydrophobic interactions, while same cavity in mutant form four hydrogen bonds (Glu154-Ala123, Glu154-Arg102, Ser155-Asp142, and Val157-Gly156) and higher hydrophobic interactions. Higher intra-atomic contacts leads to the decrease in flexibility (increase in rigidity) by **ΔS_Vib_ ENCoM: - 0.500 kcal.mol^−1^.K^−1^** (Δ Vibrational Entropy Energy between Wild-Type and Mutant). To correlate these findings with MD simulations RMSD and RMSF were analyzed to further comment on flexibility of GBRC-NCD-370.

**Figure 1:**
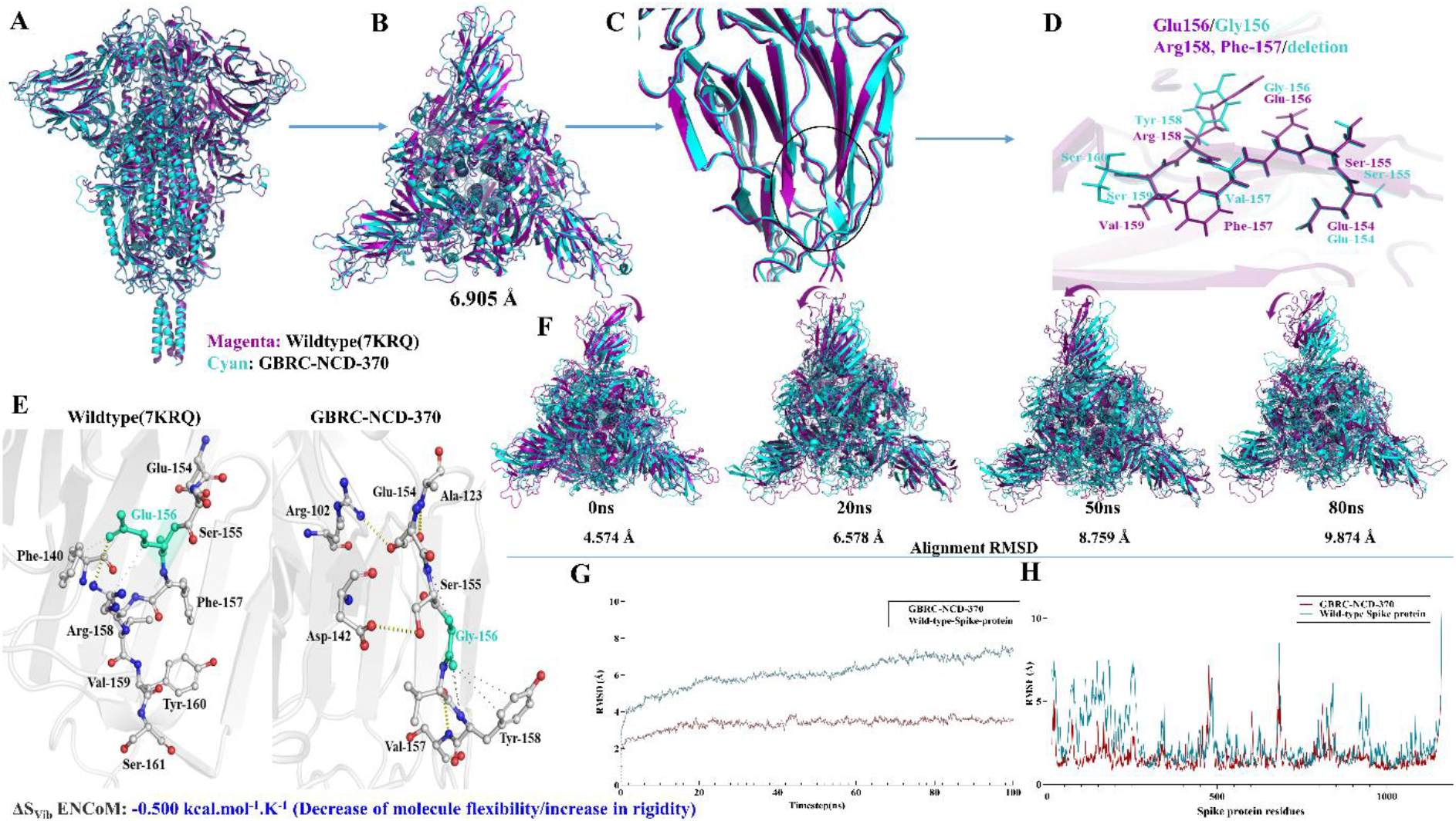
Rigidization and reduce in flexibility of N-Terminal domain of spike protein. **1A:** 3D Structural alignment of wild-type [7KQR] and GBRC-NCD-370 trimetric spike proteins with superimposition RMSD value: 6.356. Wild-type protein is shown in magenta and GBRC-NCD-370 is shown in cyan color. **1B:** Top view of trimetric spike protein. **1C & 1D**: Focusing structural difference in NTD of wild-type and GBRC-NCD-370 spike protein. **1E:** Intermolecular contacts between wild-type and mutant spike protein. **1F:** Frame superimposition of wild-type and GBRC-NCD-370 spike proteins for visualization of dynamics modes depicting difference in NTD. Magenta colored arrow showing dynamic moments of wild-type spike. **1G & 1H**: RMSD and RMSF plot generated from MD-Simulation respectively. Wild-type protein is shown in deep teal color and GBRC-NCD-370 is shown in red color.

**Table 1:**
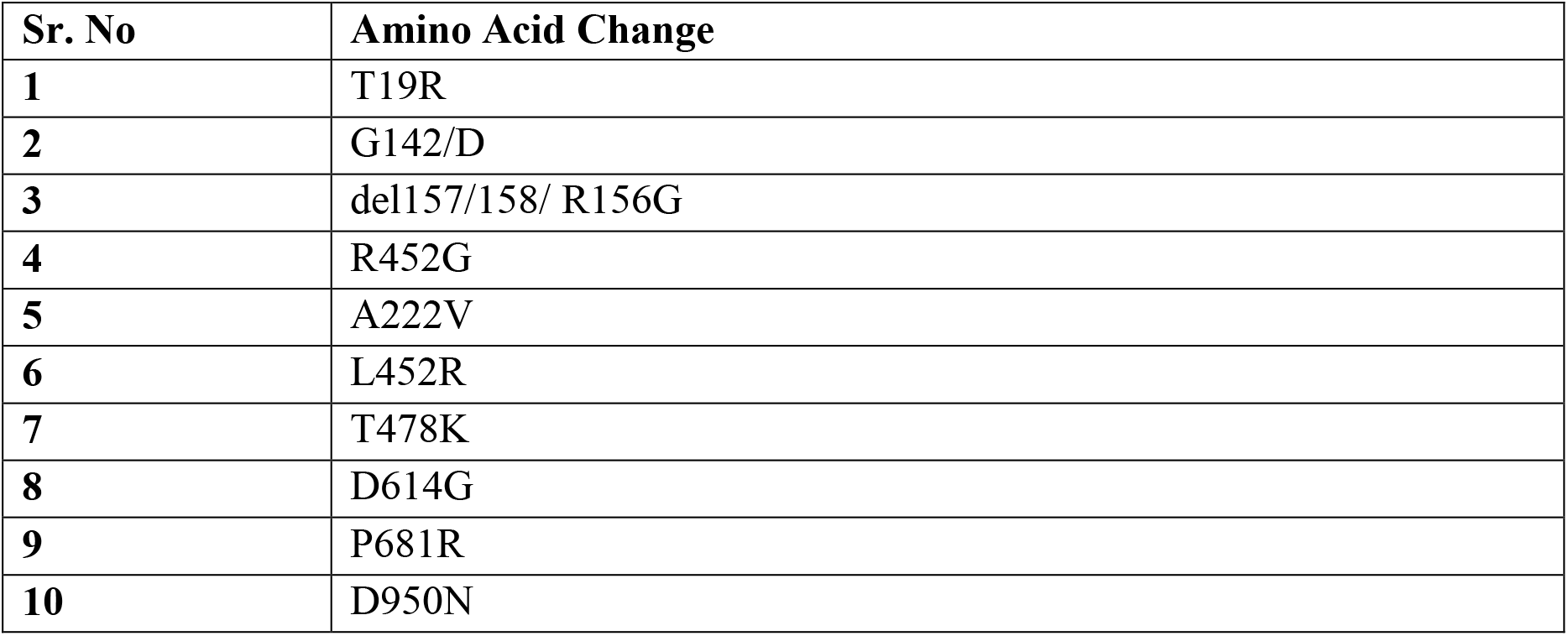
List of spike protein mutation present in GBRC-NCD-370

RMSD (Root mean square deviation) for wild-type and GBRC-NCD-370 complex observed were 5.89±0.026 and 2.54±0.018, respectively (figure1G). RMSD graph is clearly narrating that mutations in GBRC-NCD-spike protein are enhancing it stability compared to the wild-type trimeric complex (figure1H). RMSF (Root mean square fluctuation) was 3.6 Å lower in NTD of GBRC-NCD-370 spike compared to the wild-type complex. Decreased RMSF in NTD explains reduced flexibility of amino acid residues within the region. Aurélie Bornot et al 2010 precisely explains the protein flexibility in terms of RMSF and B-factors, where increasing in RMSF values is related to increased change in protein conformation. In some cases, amino acid residues are flexible though RMSF can be rigid through B-factors [37]. In our case, protein seems to be flexible in both cases through RMSF and B-factors (Supplementary figure S2). While no major change was observed with respect to other part of protein. As NTD is binding site of wide variety of monoclonal antibodies, rigidization in that region further affects the binding of antibody within both variants of spike. Principle component analysis was performed where first dominant dynamic mode PC1 among the trajectories were analyzed in VMD. Porcupines plots showing the projection of mode vectors based on the residue fluctuation throughout the trajectories were shown in Figure 2C. Length of mode-vectors in wild-type complex was higher compared to GBRC-NCD-370, which suggests that overall NTD flexibility is decreased in B.1.617.2 lineage. Increase in intermolecular contacts in mutated region further support the rigidization of GBRC-NCD-370 (figure1E).

**Figure 2:**
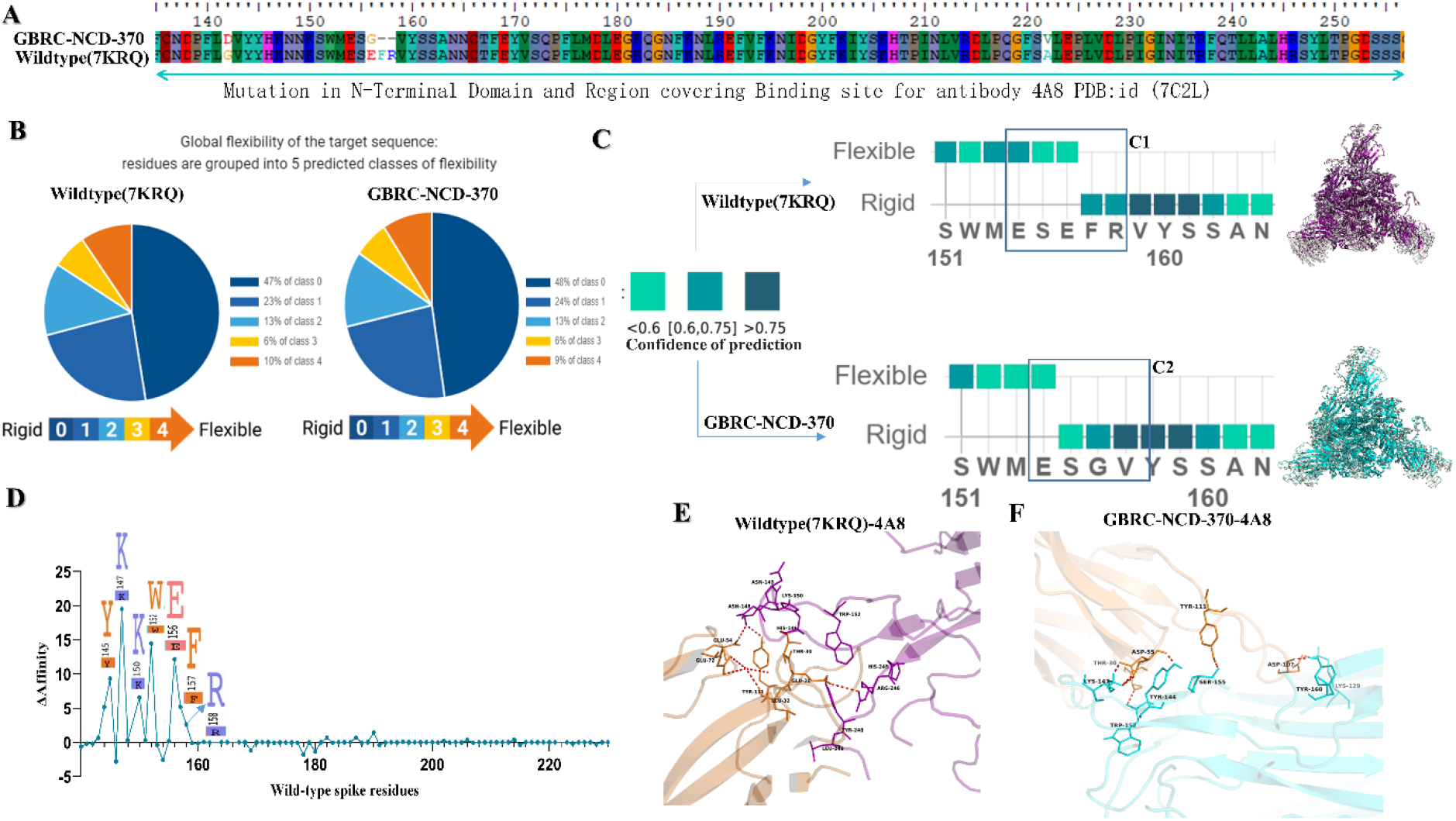
Reduce flexibility influence the binding of monoclonal antibodies (4A8) with spike protein. **2A:** Amino acid residues falling in the binding region NTD of spike. Wildtype and mutated sequences were annotated with NCBI reference/accession id number MN986947.3 and GBRC-NCD-370 respectively. **2B:** Output generated from MEDUSA to determine change rigidity and flexibility among both variants. 5 predicted classes were generated in range of 0-4, where blue region explains rigid regions and yellow to orange regions explains the flexible regions among proteins. **2C:** Flexible and rigid regions in region covering mutation. Cyan to deep teal color represents the flexible to rigid region with COP (confidence of prediction) with <0.6 and >0.75 respectively. Porcupines plots generated from PCA analysis were also supporting the same were shown red tube conformation with mode vectors. **2D:** Alanine residues scanning of wildtype-4A8 complex. Residues important in bind with monoclonal antibodies were shown in logo plot with positive binding affinity. **2E**: Binding pose of Wildtype-4A8 complex. Spike is shown in magenta color while 4A8 is shown in orange color. Residues involved in pivotal contacts like hydrogen bonds (red) were shown in ball and stick conformation with black colored labels. **2F**: Binding pose of GBRC-NCD-370-4A8 complex. Spike is shown in cyan color while 4A8 is shown in orange color. Residues involved in pivotal contacts like hydrogen bonds (red) were shown in ball and stick conformation with black colored labels.

Mutations in NTD were covering the binding domain for monoclonal antibodies (Figure 2A). PDB id 7CL2 was chosen as a wild-type complex of spike with 4A8 monoclonal antibody. GBRC-NCD-370 variant was docked with 4A8 was chosen as mutant complex. Glide energy of wild type A48 and GBRC-NCD-4A8 were −115.64 and −68.74 respectively. More negative energy score was showing enhance binding among protein-protein complex. MEDUSA five class predictions narrate the decrease in flexibility by 2% for the GBRC-NCD-370 compared to wild-type (Figure 2B & 2C). Spike flexibility and rigidity were analyzed in term of its binding with monoclonal antibodies by performing alanine residue scanning (Figure 2D). Wild-type-4A8 was processed through residue scanning with alanine mutagenesis to investigate the important residue for the binding with 4A8. Amino acid residues Y145, K147, 150K, 152W, 156E, 157F, and 158R showed positive binding affinity values with 4A8 upon mutating these residues to alanine (Figure 2D). These results clearly indicate that the mutations in the NTD domain of spike caused decrease in binding of 4A8 antibody in B.1.617.2 lineage. To further explore the effect, the mutations in NTD with respect to affinity with 4A8, MD simulations were performed in duplicates and binding energies among both variants were analyzed.

### 3.2 Effect of Arg158, Phe-157/del and E156G in N- terminal domain with respect to the host immunity

MD simulations of both mutant and wild type spikes with 4A8 antibody explored the effect of mutation Arg158, Phe-157/del and one amino acid mutation E156/G in terms binding with monoclonal antibodies and depict the case of immune evasion. RMSD of GBRC-NCD-370-4A8 and wildtype-4A8 was 20.147±0.526Å and 16.142±0.453Å, respectively (Figure 3A). In GBRC-NCD-370-4A8 platue was reached after 65 ns and jumps were observed during the MD simulations, while wildtype-4A8 was found to be stable after 20ns only. These major difference among the both trajectories shows that GBRC-NCD-370-4A8 complex is having 4 Å less RMSD, leads to more stable than wildtype-4A8. Hydrogen bonds formation within GBRC-NCD-370-4A8 complex was lower (on an average 6 hydrogen bonds difference was observed) compared to wildtype-4A8 (figure 3B). Hydrogen bond formation clearly indicates reduced interaction of antibodies in B.1.617.2 lineage.

**Figure 3:**
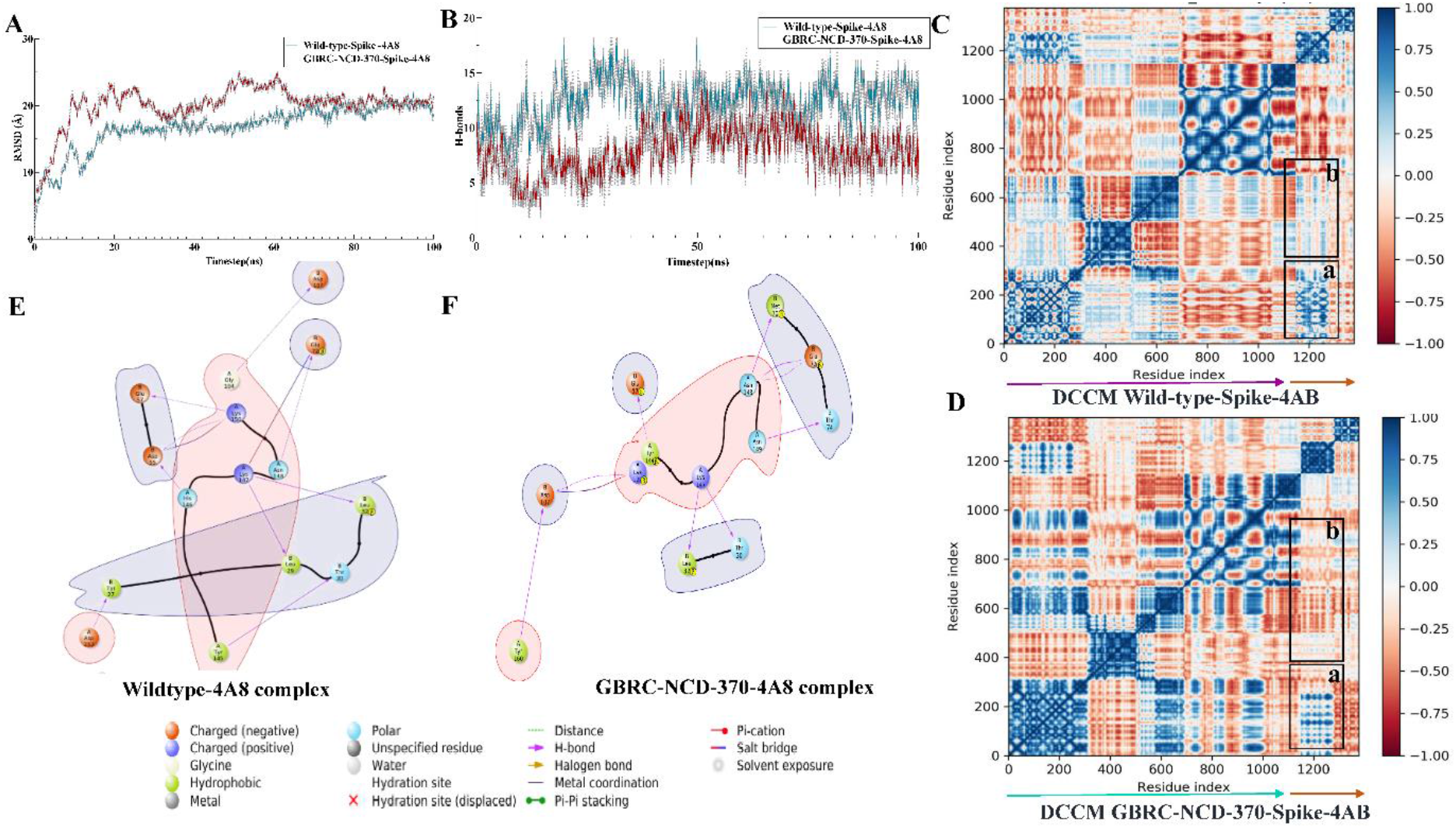
MD analysis of Spike-antibodies complexes. **3A**: RMSD (root mean square deviation) within Wildtype-4A8 (cyan) and gbrc-ncd-370-4A8 (mutant) complex. **3B:** Hydrogen bonds formation within Wildtype-4A8 (cyan) and gbrc-ncd-370-4A8 (mutant) complex. **3C**: Dynamics cross-correlation matrix obtained from trajectories analysis of wild-type-4A8 complex. Spike protein shown in magenta arrow and orange arrow is indicating 4A8. **4D:** Dynamics cross-correlation matrix obtained from trajectories analysis of GBRC-NCD-370-4A8 complex. Spike protein shown in cyan arrow and orange arrow is indicating 4A8. Blue to red color represents the Cij values between 1 to −1. No cross correlation was shown by white color. **3E & 3F:** Energy minimized structure obtained through MMGBSA for Wildtype-4A8 and GBRC-NCD-370-4A8 respectively. Positive chagrined, negative charged amino acid residues were shown in orange and blue color respectively

Binding energy among the complex was analyzed through MMGBSA. Major energies contributing to the complex formation were elucidated in table 2 with bold text. Spike-4A8 complex formation is driving through the major electrostatic, covalent, ionic-interactions, lipophilic (hydrophobic) and Vander-Waals interactions within both complexes. Overall free energy binding ΔG in wild-type and GBRC-NCD-370 complex is −119.086±19.42 and - 55.496±14.57, respectively. Interaction in energy minimized structure obtained through MMGBSA approach is shown in figure 3E & 3F. In wild-type complex overlapping strong interaction between charged negative (orange) and charge positive residues (blue) is way higher compare to mutant. For example A-Lys147: B-Glu72, A-Lys150: B-Glu57 & B-Glu55 were forming hydrogen bonds and salt bridges in spike (A) and 4A8 (B). Jason E Donald and group suggests that salt bridges were geometric specific and designable interactions [38,39]. Lys150 is forming salt bridge and hydrogen bonds with two negative charged amino acids Glu57 and Glu55 (Figure 3E & 3F). These kind of favorable interactions are formed in wild-type spike but absent in GBRC-NDC-370, leads to conclude that geometry of NTD in spike had changed as such that it is reducing the strong interaction with 4A8 in GBRC-NCD-370 (mutant) (Table:2). This kind of salt-bridges are favorable exist in hydrophobic environment [40], leads to higher lipophilic energy in wild type (−30.334 Kcal/mole) compare to GBRC-NCD-370 (−11.9757Kcal.mole). Overall Wildtype spike seems to have better binding with monoclonal antibodies compared to the GBRC-NCD-370, which leads to conclude that there is possible case of immune evasion among B.1.617.2 lineage.

**Table 2:** Differences in energy components contributing to Complex formation within Wild-type (7KRQ) and Mutated GBRC-NCD-370 monoclonal antibodies (4A8).

| Energy components | Wildtype-4A8 | GBRC-NCD-370-4A8 |
| --- | --- | --- |
| **Glide energy** | **-115.64** | **-68.74** |
| $\Delta G$ Binding | -119.0869622 | -55.4968 |
| $\Delta G$ Electrostatic energy | -791.09011 | -357.715 |
| $\Delta G$ Covalent energy | -14.19260182 | -7.50638 |
| $\Delta G$ Hbonds energy | -8.385258773 | -5.17996 |
| $\Delta G$ Lipophilic energy | -30.33398583 | -11.9757 |
| $\Delta G$ pi piinteraction energy | -2.106536604 | 1.156437 |
| $\Delta G$ selfcontactcorrelation | -0.023560517 | -0.15669 |
| $\Delta G$ Solv_GB | 781.5183498 | 434.7153 |
| $\Delta G$ vdw energy | -82.85846217 | -108.834 |

Dynamics cross-correlation matrix (DCCM) of wild-type and GBRC-NCD-370 spikes with 4A8 is shown in figure 3. In DCCM wild-type-spike-4A8 is showing higher intensity of blue color compared to the GBRC-NCD-370-spike-4A8. Positive C_ij_ values indicate blue colors leads to better interaction profile between those residues. NTD region covers 17-305 amino acid residues where major residue contributing in direct contact was shown in figure 2. In the region covering orange arrow (antibody 4A8) intensity of blue color is higher, indicating more positive cross correlation with respect to NTD region of spike wild-type compare to GBRC-NCD-370. In wild-type complex NTD residues have showing higher negative cross-correlation compare to GBRC-NCD-370 (Figure 3C &3D). Results of negative cross-correlations in some regions were completely correlating with change in flexibility of NTD. Higher intensified positive cross-correlation showed structural compactness among the NTD in GBRC-NCD-370, which can be unfavorable for the antibody binding. Overall cross-correlation among 4A8 which were shown in box a and b with respect to spike, were positive in wild-type and negative in GBRC-NCD-370 supporting the case of antibody escape.

Mutant have increased rigidity which is concluded using Insilco methods. Increase in interatomic contacts have enhanced the rigidity which is shown by two platforms pymol and Medusa. Further PCA analysis and porcupine plot were also showing the decreased in flexibility (increased rigidity). Andrey Karshikoff and group had shown binding site residues of protein have flexible tendency, to have better interaction with ligand (here the case of antibody) [41]. As binding site residues are more engaged with intra-atomic contacts itself, tendency of sharing contacts with outsider protein will be less [42]. Based on results obtained through higher RMSF and Porcupine plots it was hypothesized that rigidization among NTD leads to antibody escape.

## 4. Conclusion

The present study addressed the critical structural and genomic determinants of the SARS-CoV-2 (B.1.617.2/Delta) variant which is most dominant in India during the second wave and spreading quickly in different geographical regions of the globe. The E156G and Arg158, Phe-157/del mutations in NTD of spike protein of SARS-CoV-2 (B.1.617.2/Delta) variant showed more rigidity and reduced flexibility compared to Wuhan isolate. Further, our study showed possible case of immune escape by demonstrating reduce binding of mutant spike compared to Wuhan isolate with reported antibody known to bind NTD of spike protein thereby providing insights into the structural basis and highlight the impact of the key mutations for the higher transmissibility, pathogenicity and virulence. Therefore, it is important to better monitor and identify the new emerging variants of SARS-CoV-2 using genome sequencing and surveillance that may have increased transmission, virulence and altered antigenicity evolved over time for epidemiological significance.

## Credit authorship and contribution statement

AC performed all Insilco experiments, analysis, writing original draft preparation, data curation. DK analyzed the sequences, write and edited the manuscript. AP, MJ, CJ writing review and editing, supervision, project administration, funding acquisition

## Acknowledgement

Authors would like to acknowledge Department of Science and Technology (DST), Government of Gujarat for infrastructure support for the research work. We also thank Gujarat University for providing us GPU accelerated computer facility to enhance the speed of our work.

