## Supplemental Figures for "E156G and Arg158, Phe-157/del mutation in NTD of spike protein in B.1.617.2 lineage of SARS-CoV-2 leads to immune evasion through antibody escape" (from another version of this preprint: version 2, 2021-06-14)

**Supplementary files**

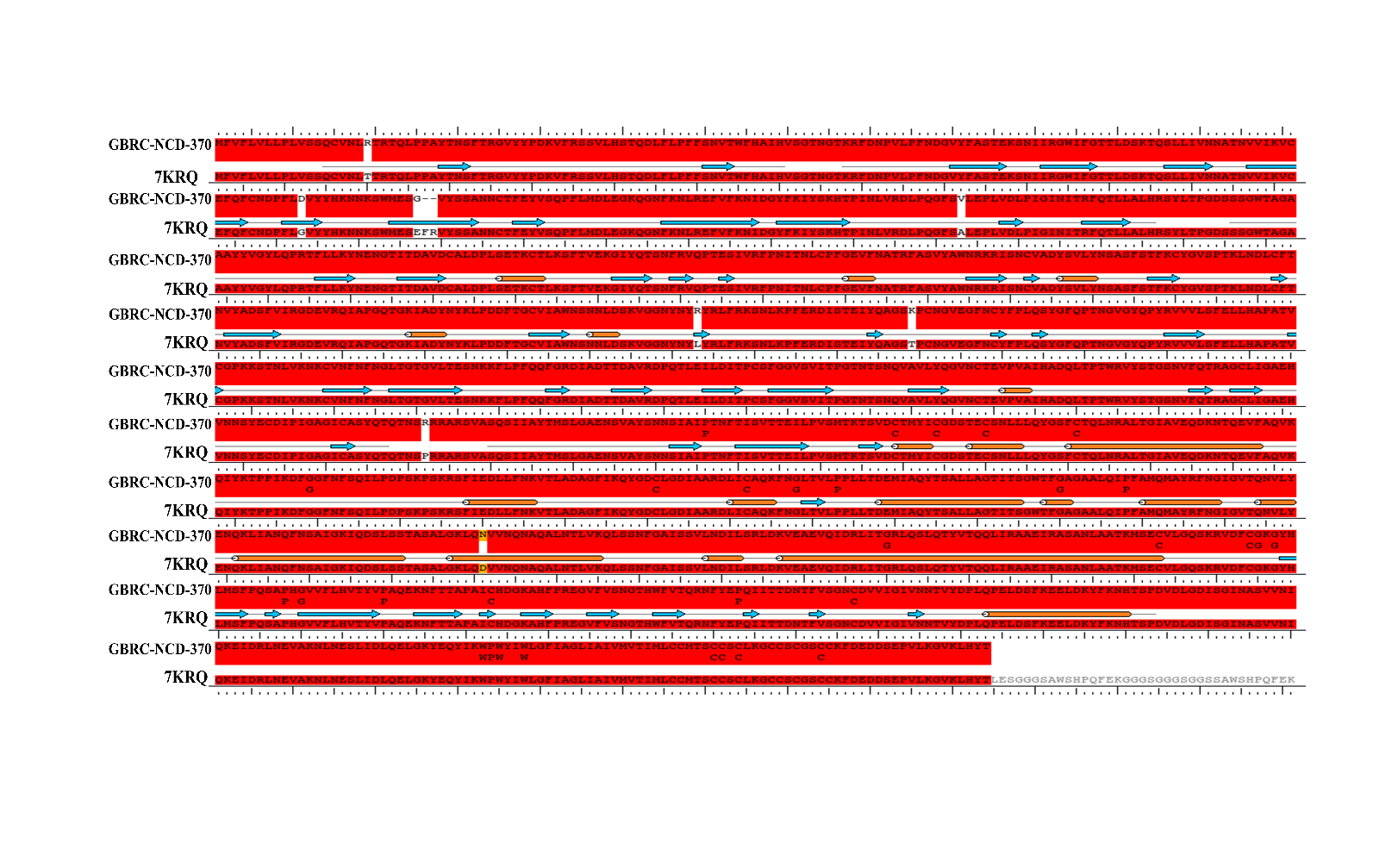

**Supplementary Figure S1:** Sequence alignment of GBRC-NCD-370 and 7KRQ. Secondary structure elements were shown in Orange (alpha helix) and blue (beta-sheets). Mutated Residues are not highlighted with red color.

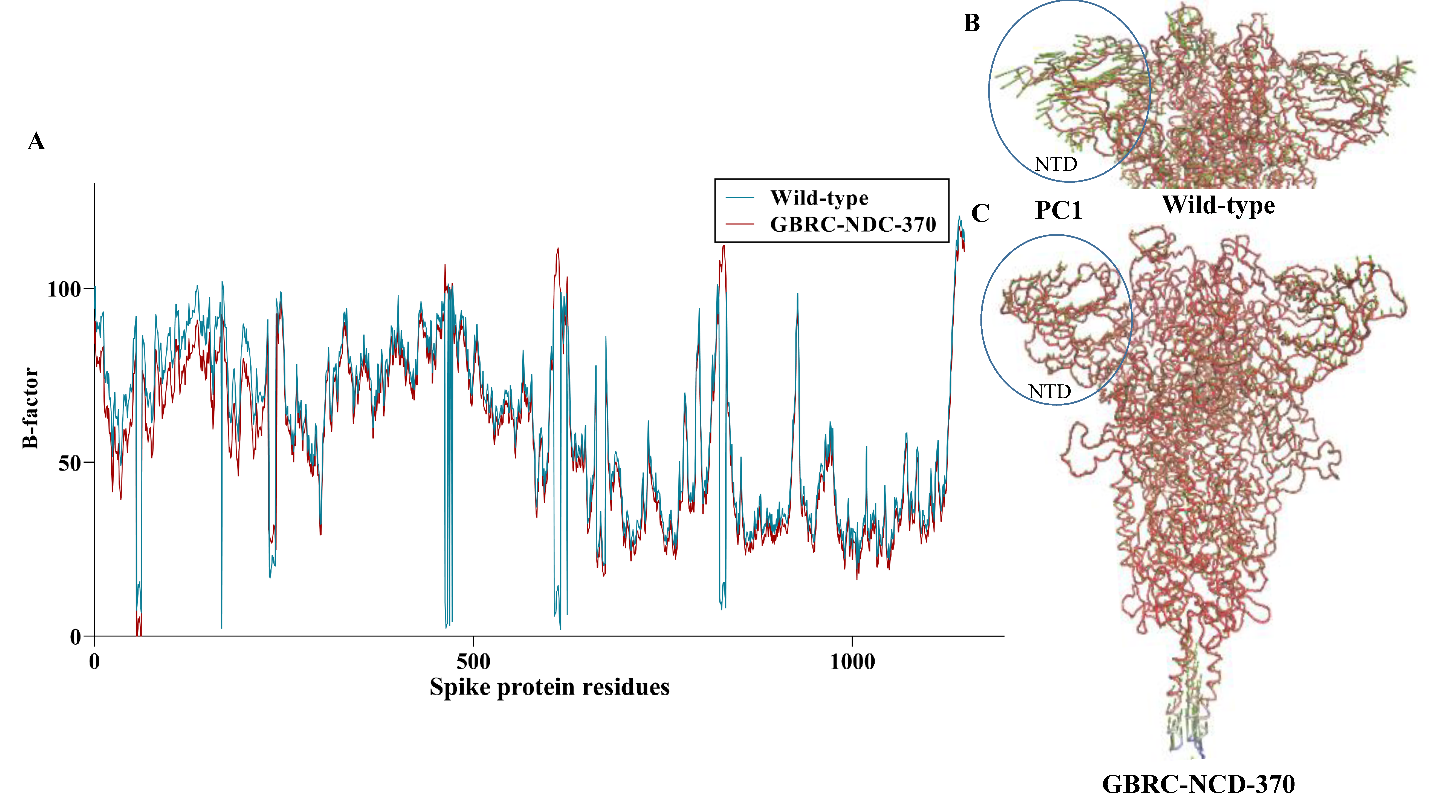

**Supplementary Figure S2:** Protein flexibility during MD simulation **(A)** B-factors among both variants of spike protein wild-type is shown in blue and mutant is shown in red. **(B)** Wild-type PC1 showing reduced flexibility in NTD of spike. **(C)** GBRC-NCD-370 PC1 showing reduced flexibility in NTD of spike.
